# Multiplexed biochemical imaging reveals caspase activation patterns underlying single cell fate

**DOI:** 10.1101/427237

**Authors:** Maximilian W. Fries, Kalina T. Haas, Suzan Ber, John Saganty, Emma K. Richardson, Ashok R. Venkitaraman, Alessandro Esposito

## Abstract

The biochemical activities underlying cell-fate decisions vary profoundly even in genetically identical cells. But such non-genetic heterogeneity remains refractory to current imaging methods, because their capacity to monitor multiple biochemical activities in single living cells over time remains limited^1^. Here, we deploy a family of newly designed GFP-like sensors (NyxBits) with fast photon-counting electronics and bespoke analytics (NyxSense) in multiplexed biochemical imaging, to define a network determining the fate of single cells exposed to the DNA-damaging drug cisplatin. By simultaneously imaging a tri-nodal network comprising the cell-death proteases Caspase-2, -3 and -9^2^, we reveal unrecognized single-cell heterogeneities in the dynamics and amplitude of caspase activation that signify survival versus cell death via necrosis or apoptosis. Non-genetic heterogeneity in the pattern of caspase activation recapitulates traits of therapy resistance previously ascribed solely to genetic causes^3,4^. Chemical inhibitors that alter these patterns can modulate in a predictable manner the phenotypic landscape of the cellular response to cisplatin. Thus, multiplexed biochemical imaging reveals cellular populations and biochemical states, invisible to other methods, underlying therapeutic responses to an anticancer drug. Our work develops widely applicable tools to monitor the dynamic activation of biochemical networks at single-cell resolution. It highlights the necessity to resolve patterns of network activation in single cells, rather than the average state of individual nodes, to define, and potentially control, mechanisms underlying cellular decisions in health and disease.

---

Genetically identical cells exhibit significant heterogeneity in their response to a stimulus that is invisible to ensemble analyses of cell populations^5,6^. Recent studies provide multiple lines of evidence that non-genetic heterogeneity is a major determinant of physiology and pathology in multicellular systems^7–10^. Analysis of the mechanisms underlying non-genetic heterogeneity relies on the ability to observe biochemical reactions in intact single cells^5,11–13^, for which new microscopy approaches have been developed^1,12,14^. However, available methods remain limited in their capacity to monitor multiple biochemical activities in single living cells over time, challenging our understanding of how the dynamic response of biochemical networks can elicit heterogenous cellular phenotypes.

This challenge is exemplified by the cell-fate decisions that follow the exposure of normal or diseased cells to DNA damage, which culminate variously in damage repair, survival or death^5,6,11,15,16^. Ensemble analyses implicating the cell-death activating proteases Caspase-2, -3 and -9 in the mechanisms that dictate cell fate after DNA damage suggest that the apical activation of either Caspase-2 or -9 is instrumental^4,17–22^. In particular, Caspase-9 activation is proposed^3,4,17,23^ to determine the therapeutic sensitivity of cancer cells to the genotoxic drug, cisplatin, which is widely used in the treatment of ovarian, lung and other common cancers^24^. Although there is growing evidence that non-genetic heterogeneity is a significant cause of therapeutic resistance to anticancer drugs including cisplatin^7,8^, underlying mechanisms that may open avenues to predict and potentially overcome resistance remain poorly characterized and difficult to model.

Here, we report the development of a unique approach to monitor over time small biochemical networks in single living cells based on pairs of GFP-like fluorescent and non-fluorescent proteins (NyxBits), fast electronics and computational tools (NyxSense) of new design, a technology platform we will make available to the community. The NyxBits are a family of bright (reporter) and dark (quencher) GFP-like proteins that can be rationally combined to generate genetically encoded sensors for biochemical multiplexing. The NyxBits reporters are bright fluorophores—mTagBFP^25^, mAmetrine^26^ and mKeima^27^—that are excited with a single laser (400-440nm) and utilize the whole visible spectrum efficiently (**Fig. 1** and **Extended Data Fig. 1**). The NyxBits quenchers are chromoproteins or fluorophores with very low emission—sREACh^28,29^, *ms*CP576 and tdNirFP—that attain high FRET efficiencies, yet minimise spectral cross-talk between the FRET pairs. To develop this system, we undertook a laborious analysis of tens of donor-acceptor pairings to provide a suitable multiplexing platform, minimizing aggregation and cross-talk of different reporters, as briefly described in **Supp. Note 1** and **Supp. Tables 1-2**. NyxBits were thus utilized to sense VDTTDase, DEVDRase and LEHDase activities as probes for the surrogate activities of Caspase-2 (blue), Caspase-3 (green) and Caspase-9 (red), respectively. NyxBits-based caspase sensors respond as expected and with an excellent dynamic range upon treatment of HeLa cells with 4 μM STS for 8 hours; a response that is significantly suppressed, as expected, by well-characterized inhibitors (**Extended Data Fig. 1**).

**Figure 1.**
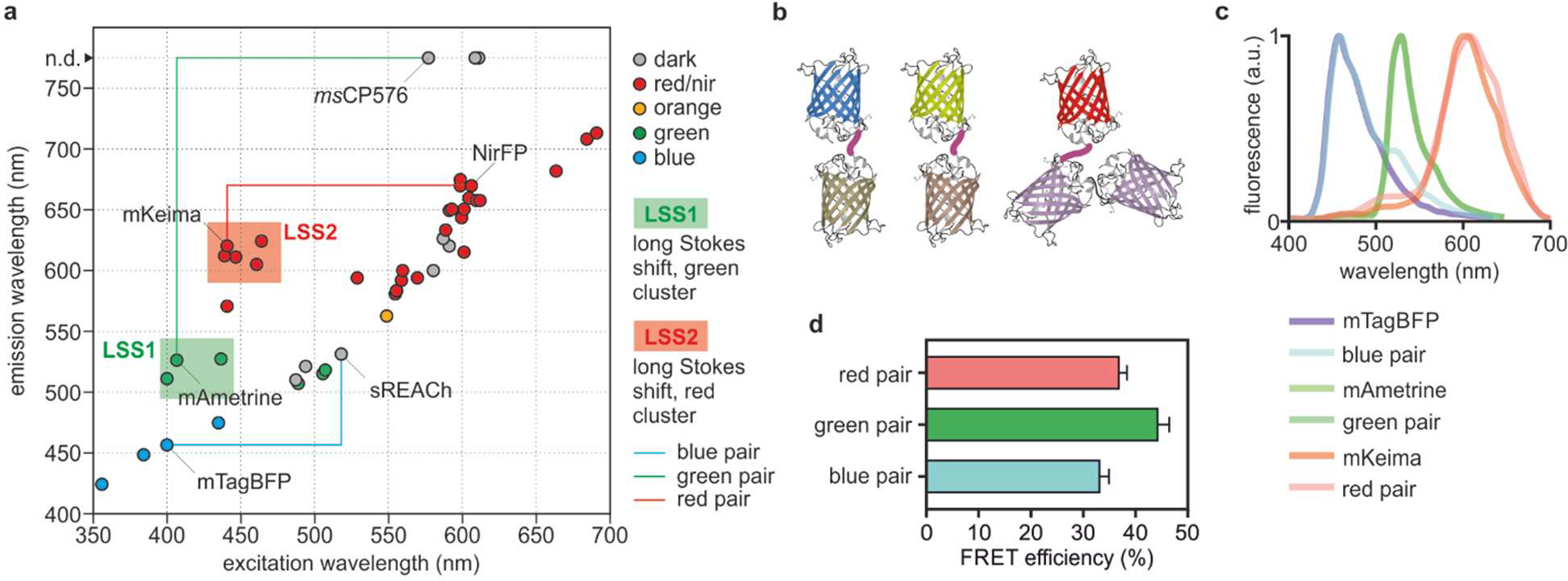
Multiplexing biochemical networks in single cells. We have tested optimal pairing of bright donor fluorophores at varying Stokes shifts with dim or dark acceptors to engineer a biochemical multiplexing platform based on FRET sensors (**a**, see also **Supp. Tables 1** and **2**). Three FRET pairs (mTagBFP:sREACh, mAmetrine:*ms*CP576 and mKeima:tdNirFP) for biochemical multiplexing (**b**) can be readily measured by multi-colour FLIM. Donor fluorophores (**c**, saturated colours) and FRET pairs (**c**, unsaturated colours) are efficiently distributed over the visible spectrum and exhibit high FRET efficiencies (**d**).

We thus generated a single plasmid encoding a poly-protein carrying the three caspase sensors, and engineered a monoclonal stably-transfected HeLa cell line (Nyx.C239). Nyx.C239 cells permitted us to study, for the first time, the dynamic activity of the caspase network in response to Cisplatin. Parental and Nyx.C239 cells exhibit similar responses both in the processing of caspases and inhibitor-of-apoptosis (IAP) family proteins demonstrating the minimal invasiveness of the methodology (**Extended Data Fig. 2**). NyxBits sensors can be quantified only by Fluorescence Lifetime Imaging Microscopy (FLIM)^14,28^. To overcome common limitations in speed and dynamic range of FLIM, we concurrently developed fast photon-counting electronics that have become commercially available very recently (**Supp. Note 2**). Moreover, we developed the open source NyxSense (**Supp. Note 3**) software that implements *ad hoc* data analysis algorithms based on multi-dimensional phasor^30–32^ fingerprinting. NyxSense enables quantitative unmixing of biochemical information from the NyxBits platform and multi-colour FLIM, delivering an unprecedented level of multiplexing. The combination of these innovations permitted us to measure directly in single living cells, the dynamic and the heterogeneous activity of multiple biochemical reactions over time (**Fig. 2d-g**, **Extended Data Fig. 3c-d**, **Supp. Videos 1 and 2**) over thousands of phenotypically annotated cells

**Figure 2.**
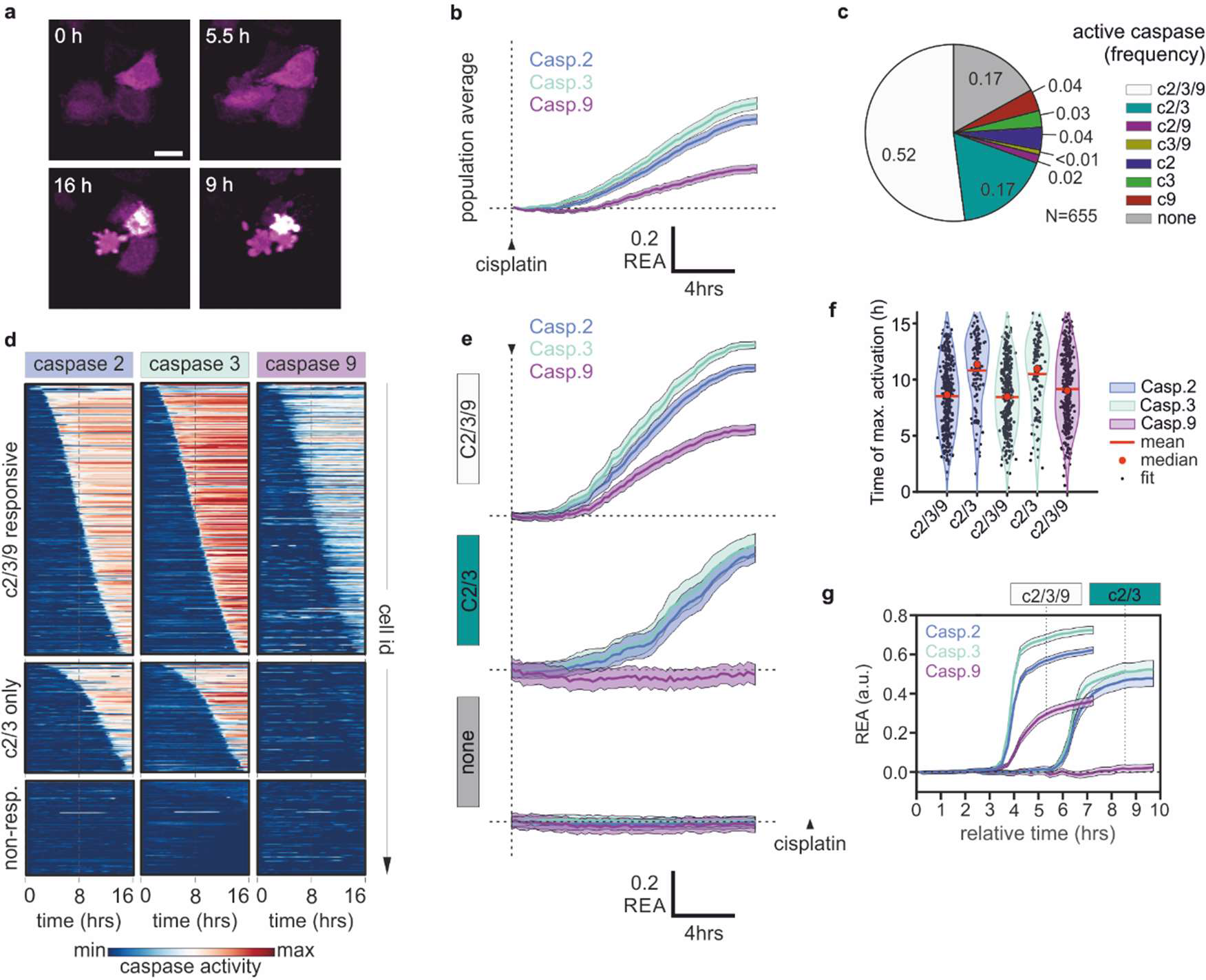
Deconvolution of dynamics and heterogeneity of biochemical networks. Time-lapse biochemical imaging (**a**) is thus enabled by FRET sensors utilizing these novel pairs, *e.g.* as Caspase-2, -3 and -9 sensors. From typical populations averages (**b**, bootstrapped averages and 95% intervals shown), it is possible to deconvolve distinct cellular populations that utilize different patterns of caspase activations (**c**). Single-cell traces can be then analysed individually (**d**) or as averages representative of each population (**e**). Moreover, the timing of caspase activation (**f**) can be utilized to synchronize all traces within each population, to determine the prototypical responses as if all cells responded homogeneously and synchronously (**g**), information typically lost during ensemble measurements. In the C239-responsive population, the average cleavage of Caspase-2 and -3 substrates appear simultaneous, while a robust cleavage of the Caspase-9 sensor is detected ~40 minutes later (one tail t-test, p<10-6, N=331). The C23-only population exhibits also a delayed cleavage of Caspase-2 sensor for ~20 minutes (p<0.05, N=101).

The large majority (~80%) of Nyx.C239 cells trigger cell death and activate caspases on average after ~9.1±0.2 hours (mean and standard error, N=560; **Fig. 2**) consistent with data acquired on parental cells (**Extended Data Figs. 5i** and **2a**). But single-cell biochemical traces revealed the highly heterogeneous activation of the caspase network (**Fig. 2c-d**) that is concealed in these population averages (**Fig. 2b** and **Extended Data Fig. 2**). The time at which each cell triggers cell death is highly variable^6,8^ (**Fig. 2d,f**). Moreover, multiplexing of caspases in single living cells also exposes a previously unreported variability in patterns of caspase network activation. About 50% of cells activate all three caspases (C239-responsive, N=331) at 8.4±0.2 hours on average, but ~20% of the population trigger Caspase-2 and -3 robustly only two hours later (10.3±0.3, N=101; **Fig. 2d-g**), without any apparent activation of Caspase-9 (C23-only). A minority (<14%) of cells activates different pairs of caspases or even individual caspases (**Fig. 2c** and **Extended Data Fig. 4d**), providing additional evidence for the specificity of each sensor.

Using the time of half-maximal cleavage of the Caspase-3 sensor as a landmark to annotate activation patterns, we were able to quantitively describe such an asynchronous and heterogeneous process. Analysed in this way, the sharp activation of caspases at single-cell resolution contrasts with the shallow responses measured in population averages. Moreover, the analysis unambiguously reconstructs the distinct response of C239-responsive and C23-only populations (**Fig. 2d,g**). On average, cells that trigger Caspase-9 together with Caspase- 2 and -3 die earlier, more promptly cleave Caspase-2 substrate, and exhibit a steeper slope in the activation of all three caspases (**Extended Data Fig. 4b**).

A series of further experiments were performed to extensively validate our observations. First, we excluded that the C23-only sub-population was an artefact of the particular clone of Nyx.C239 we used, by confirming the results via transient transfection of a dual Caspase sensor as shown in **Extended Data Fig. 4e-g**. Data shown in **Extended Data Fig. 4e** also confirms that the different dynamics of Caspase-9 is not caused by the use of the red NyxBits as similar traces are acquired with the green NyxBits. Second, we excluded that our observations represent a cell-cycle dependent effect, by repeating the experiments in G1- and G2-synchronized Nyx.C239 cells without observing differences (**Extended Data Fig. 4h-j**). Third, we excluded that the Caspase-9 negative/Caspase-3 positive population was an artefact of intensive multiplexing, through flow cytometric analysis of cleaved PARP and active Caspase-9 (**Extended Data Fig. 5a-b**). Flow cytometry analysis of active Caspase-9 does not exhibit a high dynamic range, and when both time-lapse single cell traces are analysed by gating like flow data, the prevalence of the Caspase-9 positive and negative populations are consistent across the orthogonal assays. Thus, taken together, these results validate our finding that a profound cell-to-cell variability in the pattern of activation of the caspase network determines the time of death of individual cells.

We then classified different phenotypes (**Fig. 3a-b; Supp. Video 1** and **2**) to distinguish mechanisms of cell death after cisplatin-induced DNA damage. We identified first, a typical apoptotic phenotype (phenotype (PT)1, ~70%) characterized by shrunken cells; second, cells exhibiting an incomplete and often slower shrinkage accompanied by blebbing (PT2, ~5%); and third, cells that exhibit only plasma membrane blebbing (PT3, ~5%) with no apparent loss of volume. Independent experiments demonstrated that PT1 cells are apoptotic, while PT2 and PT3 correspond to necrotic cells (**Extended Data Fig. 5c-e**). Notably PT2/PT3 necrotic cells die significantly later than PT1 apoptotic cells (12.5±0.4 hours N=58 *versus* 10.3±0.2 hours N=459) both in HeLa (**Extended Data Fig. 5i**) and Nyx.C239 (**Extended Data Fig. 4a-c**). A ~2 hours difference in death phenotypes is reminiscent of the delay between the activation of the caspase network in the C239-responsive and C23-only populations. Thus, we hypothesised that the C23-only population represents the necrotic fraction of Cisplatin-induced cell death. Cross-tabulation of caspase activities with the death phenotypes PT1-3 (**Fig. 3c**) suggests patterns of activity are indeed associated with apoptosis or necrosis. The C23-only population is more frequent in PT3 cells (**Fig. 3c**) and PT3 cells exhibit a more modest cleavage of the LEHD substrate compared to PT1 cells (12±4% *versus* 27±1%, **Fig. 3d**). Thus, it is apparent that those cells that exhibit a robust activation of all caspases, die earlier and are more likely to trigger apoptosis, whereas a small but significant fraction of cells exhibit shallower caspase activation, lower Caspase-9 activities, and a delayed cell death by necrosis. Interestingly, non-genetic heterogeneity in the pattern of caspase activation recapitulates traits of therapy resistance previously ascribed solely to genetic causes^3,4^.

**Figure 3.**
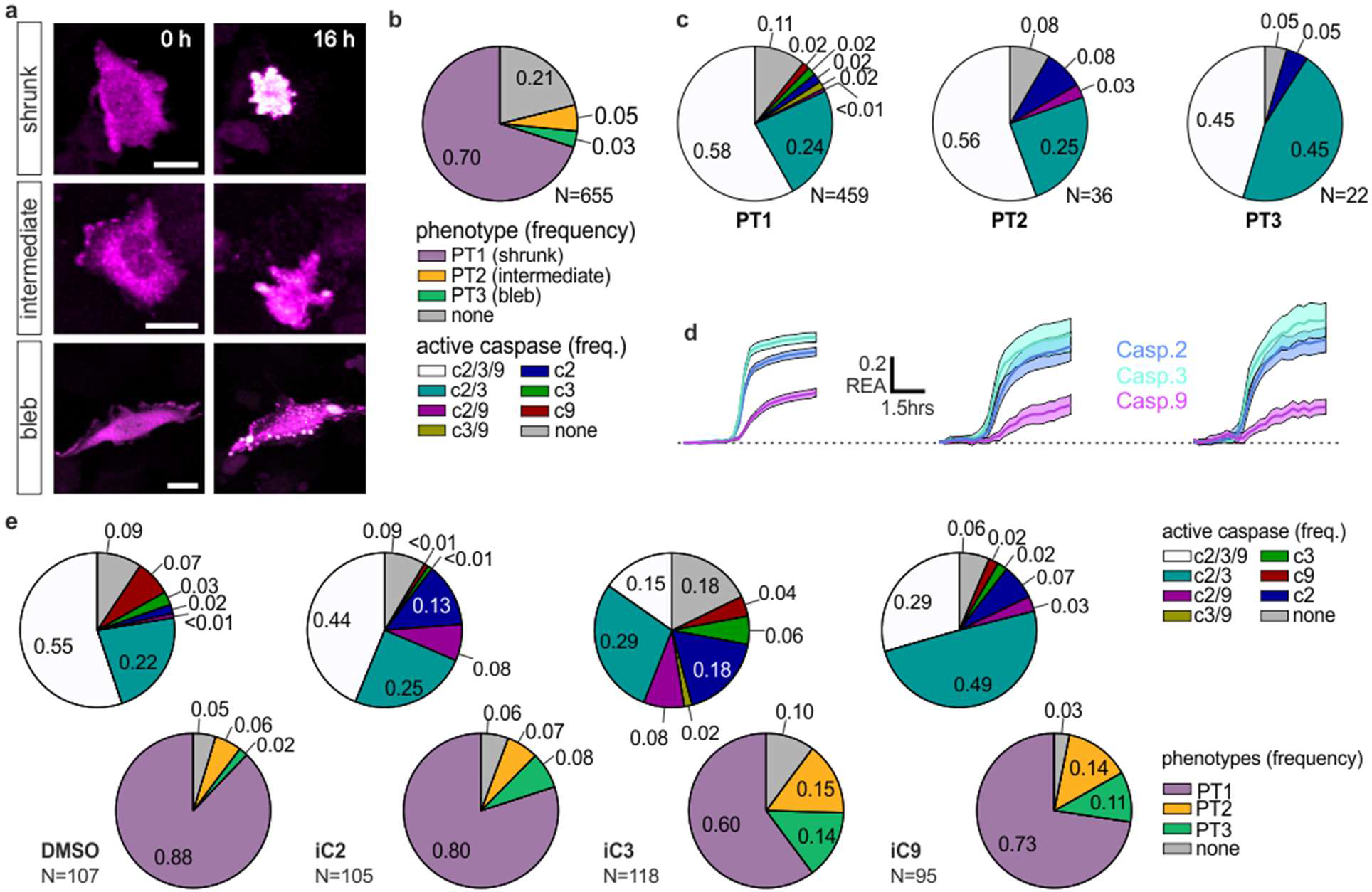
Deconvolution of phenotypes. (**a**) Cisplatin-induced DNA damage consistently causes ~80% of cells to trigger cell death exhibiting three distinct phenotypes - sudden shrinking (PT1), plasma membrane blebbing only (PT2), or partial and slow shrinkage plus blebbing (PT3). Separate experiments (**Extended Data Fig. 5**) confirm that PT1 represents apoptotic cells, while PT2 and PT3 comprise necrotic cells. (**b**) shows the frequency of PT1, PT2 and PT3. Cross-tabulation of biochemical and death phenotypes (**c**) shows that PT1 cells (apoptotic) are more likely to activate Caspase-9 than PT3 cells (necrotic). This is also evident in the lower LEHDase activity exhibited, on average, by PT3 cells (**d**). Both biochemical and death phenotypes can be altered in a predictable manner by exposure to irreversible inhibitors (**e**) for Caspase-2 (iC2: z-VDVAD-fmk) -3 (iC3: **z**-DEVD-fmk) and -9 (iC9: **z**-LEHD-fmk).

Irreversible inhibitors of Caspase-2, -3 and -9 (**Fig. 3e** and **Extended Data Fig. 6**) further elucidate the role of individual caspases in cell-fate decisions induced by the DNA damage response (DDR). First, we find that Caspase-2 inhibition reduces the fraction of C239-responsive cells. Moreover, it increases (from ~2% to ~13%) the fraction of cells that exhibit only VDTTDase activity, albeit with a rather low maximal level of cleavage (from >50% to <20%, **Extended Data Fig. 6d**) and slower kinetics. Together, these observations suggest that DDR-induced activation of the Caspase-2, -3 and -9 network cannot proceed when Caspase-2 is suppressed, despite the lower residual Caspase-2 activity that persists even after inhibitor exposure. Second, Caspase-3 inhibition causes a pronounced reduction in the frequency of the C239-responsive phenotype (from ~55% to ~15%) and apoptosis (from ~90% to ~60%). Interestingly, it increases the fraction of cells that activate only Caspase-2, thus revealing a positive feedback mechanism that connects Caspase-2 and -3. Notably, Caspase-3 inhibition also uncovers a small fraction of cells that exhibit only Caspase-2 or -9 (>10% in total), further validating the specificity of the sensors in this experimental setting. Finally, Caspase-9 inhibition induces a significant and predictable shift from the C239-responsive population (~55% to ~30%) to the C23-only population (from ~20% to ~40%).

All three inhibitors resulted in a net decrease of all caspase activities and a net delay in cell death, but a relative increase in the frequency of necrotic phenotypes PT2 and PT3 (**Extended Data Fig. 6**). Collectively, these results reveal that positive feedback circuits between caspases, either direct or indirect, set the timing of cell death. The shift from apoptotic to necrotic cells at later times after DDR induction (**Fig. 3e**) suggests an opportunity to modulate DDR-induced cell-death outcomes using clinically relevant inhibitors.

Our findings indicate that perturbation of caspase responses could change in a predictable manner the prevalence of the different death phenotypes induced by DNA damage. Therefore, we tested LCL161, a potent inhibitor of IAPs (*i.e*., a family of inhibitors-of-apoptosis proteins). IAPs restrain caspases and can be upregulated or downregulated by feedback mechanisms to either further restrain or unleash caspase activities. Accordingly, we tested c-IAP1, c-IAP2 and XIAP (also known as BIRC2, BIRC3 and BIRC4, respectively), IAPs that have a demonstrated role in DDR^12^ and that are expressed in HeLa cells. In HeLa parental and Nyx.C239 cells, c-IAP1/2 and XIAP are downregulated after treatment with Cisplatin to facilitate cell death (**Extended Data Fig. 2b**). Paradoxically, non-lethal doses of LCL161 delayed the activation of caspases and cell death by 1-1.5 hours in C239-reponsive and apoptotic cells (p<10^-3^, **Fig. 4a-c** and **Extended Data Fig.7**). However, as hypothesized, a delayed cell death resulted in increased necrosis. At the same time, we observed an increased prevalence of the C23-only population (from ~20% to ~30%) with a correspondent decrease in the C239-responsive population (from ~50% to ~35%). Therefore, these observations support the notion that non-genetic heterogeneity in the propensity to activate Caspase-9 during DDR determines the timing of the caspase-network activation eventually resulting in different cell death phenotypes.

**Figure 4.**
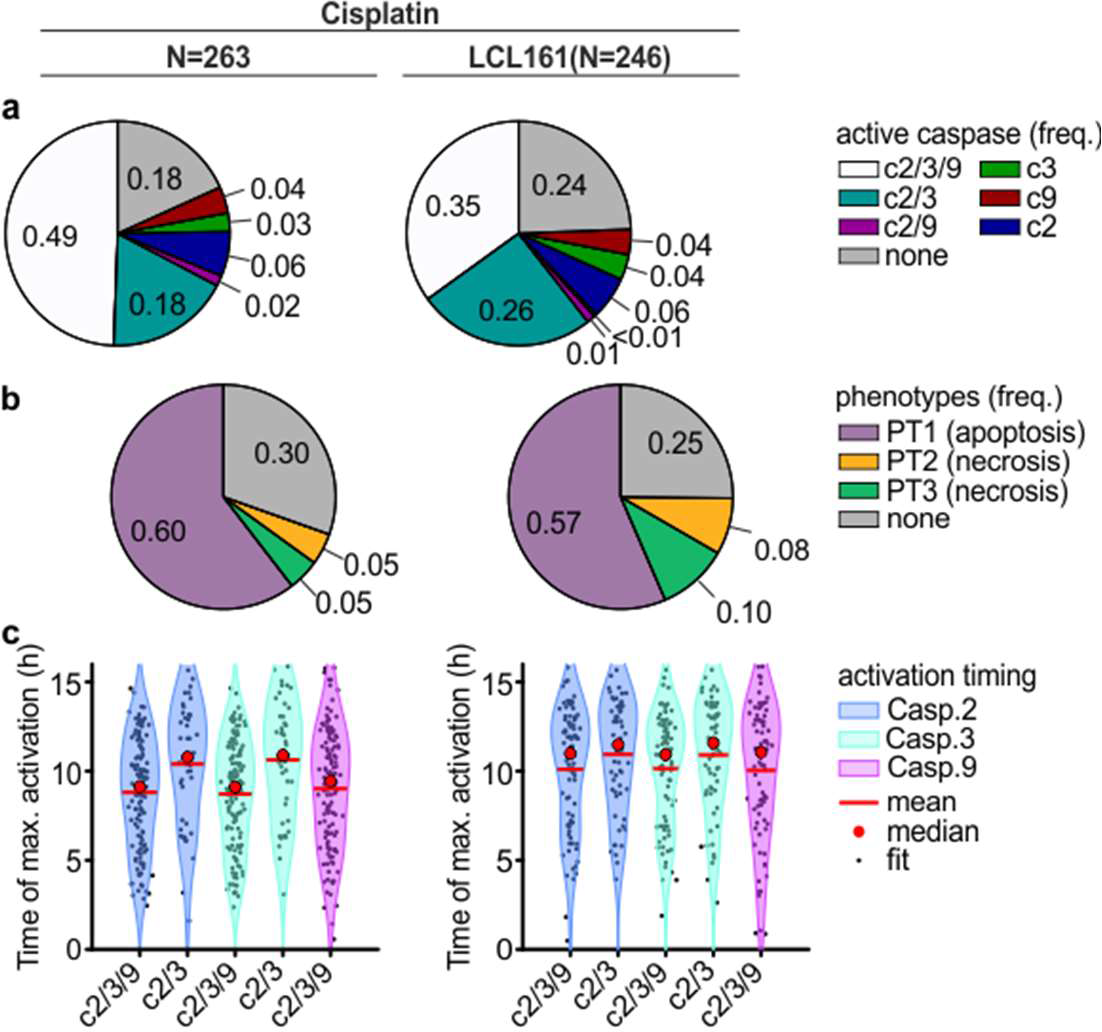
Perturbation of biochemical networks and phenotypes. IAP inhibition with sub-lethal 1 μM LCL161 alters caspase network dynamics (**a**), frequency of type of cell death (**b**) and timing (**c**) in response to Cisplatin (see also **Supp. Fig. 7**) compared to the DMSO matched control.

Cisplatin-induced depletion of ATP has been reported to be a possible discriminant between apoptosis and necrosis in response to Cisplatin treatment^33–36^. We confirmed a cisplatin-dependent steady loss of ATP (**Fig. 5a**) with the ratiometric ATP sensor ATeam^37^. Once single-cell traces are synchronized, the steep ATP-depletion measured at the population level seems, in fact, only caused by the stochastic occurrence of cell death that, both in necrosis and apoptosis, results in fast ATP loss (**Fig. 5b**). Notably, treated but surviving cells also exhibit a decrease in ATP levels (**Extended Data Fig. 8**) over time. Necrotic cells, that on average die at later times, exhibit lower ATP levels before death and a lower ATP reduction during death. These observations suggest that lower ATP levels increase the likelihood of cell death by necrosis and demonstrate a complex interaction between metabolic and signalling networks that efficiently results in clearance of damaged cells, but where cell-to-cell variability determines the type of cell death.

**Figure 5.**
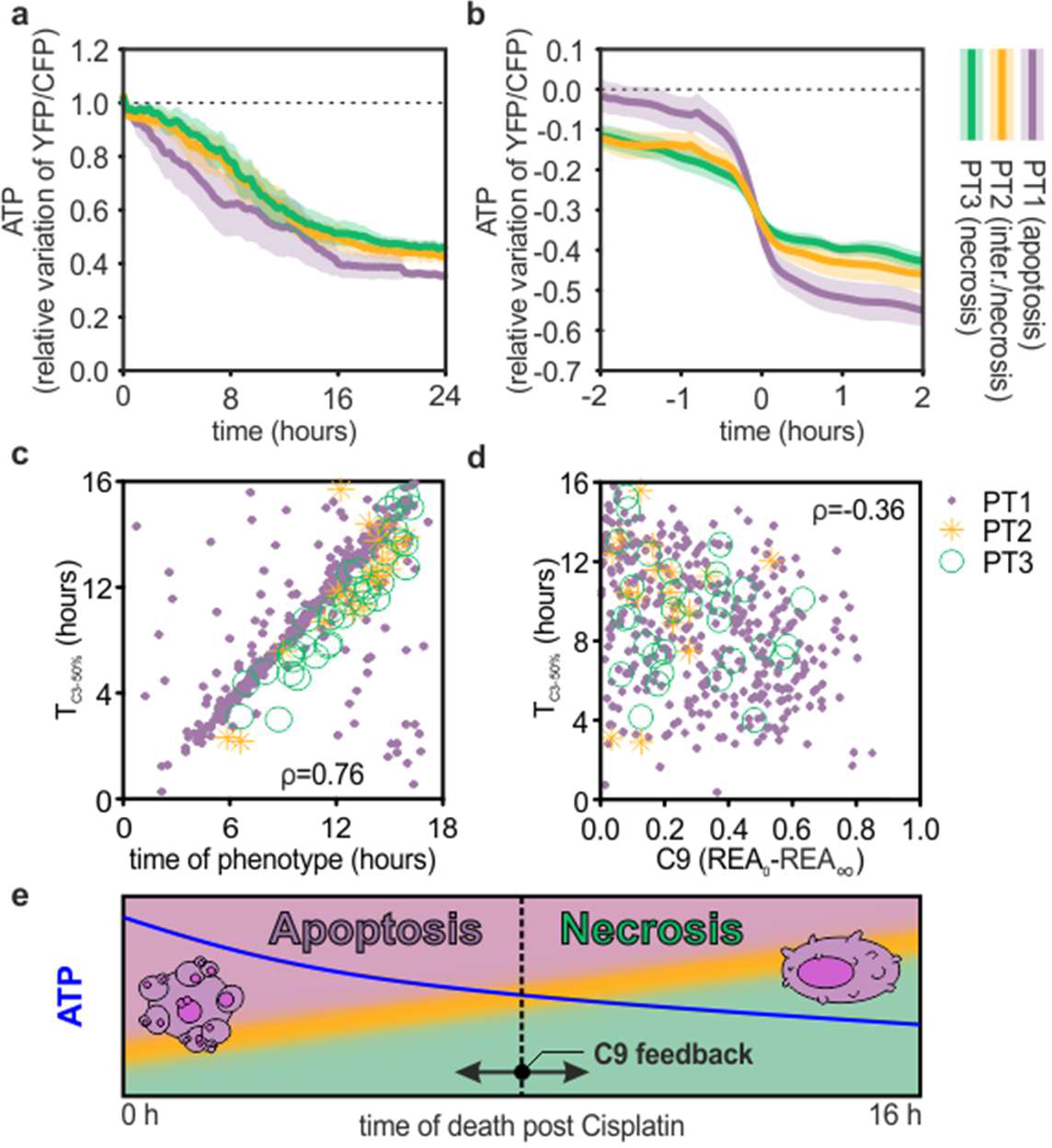
Cell-to-cell variability of biochemical networks and phenotypes. Cisplatin-induced depletion of ATP (**a**; average and 95% confidence intervals for ATeam FRET sensor) might be a determinant of cell death type (apoptosis *vs* necrosis). Single-cell traces (FRET ratio variations compared to initial values, synchronised to 30% loss) confirms differences between apoptotic and necrotic cells, but not ATP-depletion as a discriminant between the phenotypes. The prevalence of apoptosis and necrosis changes with the time after treatment (**c**) which in turn depends on the robust activation of Caspase-9 (**d**). The Spearman correlation coefficient (ρ) is shown irrespective of phenotype (otherwise ρ is equal to 0.75, 0.96 and 0.72 for the phenotypes PT1, PT2 and PT3, respectively, in **c**, or -0.36, -0.29 and -0.4 in **d**). (**e**) A hypothetical model explaining how cell-to-cell variability in caspase activation patterns over time can heterogeneously induce apoptosis or necrosis. Upon addition of cisplatin, each cell triggers a complex network of caspases (**Extended Data Fig. 8c-d**), where Caspase-2 exhibits a leading role as initiation caspase and Caspase-3 as executioner caspase, but with Caspase-9 exhibiting a critical role to provide a faster response which facilitates apoptosis. Cells with no or low activation of Caspase-9 exhibit a delayed cell death that increases the probability of necrosis, particularly in those cells with lower ATP concentration. The purple, orange and green colours depict the phenotypes PT1 to 3, respectively.

Our findings have two important implications. First, in contrast to conflicting models reported in the literature (**Extended Data Fig. 8**) that attribute apical roles to either Caspase-2 or Caspase-9 in cell-death outcomes after DNA damage^4,17–22^, our single-cell analyses reveal that the overall pattern and kinetics of the activation of a tri-nodal network comprising Caspase−2, -3 and -9 dictates cell-fate decisions in this setting (**Fig. 5e** and **Extended Data. Fig. 8c-d**). Our findings further show that robust activation of the caspase network accompanies apoptosis in the majority of cells. But a significant fraction of damaged cells exhibits less rapid, or less complete, Caspase-9 activation, inducing delayed cell death via necrosis, particularly in cells with lower ATP concentrations. Our findings exemplify approaches to manipulate the determinants of cell death in response to chemotherapeutic agents, which may improve the success of anti-cancer therapies, for instance by stimulating immunogenic cell death or averting resistance^38–40^.

Second, our work emphasizes the necessity to resolve patterns of biochemical activity in single cells, rather than through ensemble measurements, to reveal cell populations and biochemical states underlying cell-fate decisions. The unique sensing platform comprising NyxBits and NyxSense elements that we describe here enables a hitherto inaccessible application of imaging technology to achieve such resolution, and has the potential for wide applicability in the analysis of non-genetic factors important to the aetiology and treatment of many diseases through a deeper understanding of biochemical networks^38,40^, phenotypes and their variability^41,42^.

## Acknowledgements

We acknowledge funding from the Medical Research Council core grants (MC_UU_12022/1 and MC_UU_12022/8), the Wellcome Trust (090340/Z/09/Z) and the EPSRC (EP/F044011/1 and /2). We would like to thank Steve Scotcher, Howard Andrews, Phil Heard, Dave Cattermole and Martin Kyte from the mechanical and electronics workshop at the MRC LMB for their invaluable help with the engineering of our instrumentation. We also would like to thank Bryn Hardwick, Meredith Roberts-Thomson and David Perera for their initial help with cloning for some of the FRET pairs mentioned in this work, and Marina Popleteeva and Callum Campbell for discussion and support. We would like also to thank Surface Concept GmbH, Leica Microsystems Ltd and Axel Bergmann at Becker&Hickl GmbH for their assistance in integrating electronics and microscopy tools. We thank also Prof. Atsushi Miyawaki (RIKEN Brain Science Institute, Japan) for sharing the plasmids pRSET-cjBlue, pRSET-mKeima and pRSET-msCP576 (formerly named clone #20), Prof. Michael Lin (Stanford University Departments of Pediatrics and Bioengineering, USA) for pcDNA3-mNeptune, Prof. Zhihong Zhang (Britton Chance Center for Biomedical Photonics, P. R. China) for the cDNA encoding mBeRFP, Prof. Vladislav Verkhusha (Albert Einstein College of Medicine, USA) for pmTagBFP-C1, and Prof. Mark Prescott (Monash University, Australia) for help with the Ultramarine cloning. We would like also to thank Dr. Christian Frezza, at the MRC CU, for the critical reading of the manuscript.

## Methods

### Reagents

All reagents are described in **Supp. Table 6**; oligonucleotides, template sequences and cloning strategies for each sensor are described in **Supp. Tables 3** and **4** and **Supp. Methods**, respectively. All plasmids will be available through the public repository AddGene.

### Cell culture

HeLa cells (CCL 2, European Collection of Cell Cultures #93021013) were maintained in DMEM (Gibco) supplemented with 10% FCS at 37°C and 5% CO_2_ in humified atmosphere and passaged by trypsinization (Trypsin-EDTA; Gibco). HeLa cell lines stably expressing the BAK co-expression systems (HeLa-C239) were transfected with JetPRIME (Polyplus) at ~50% confluency according to the manufacturer instructions and selected in 800 ng/ml G418 (Invitrogen) for one month. A monoclonal cell line was then derived by single-cell FACS sorting and maintained under minimal selection (200 ng/ml G418).

### Western Blot

Cells were harvested lysed in 50 μl of RIPA buffer (50 mM Tris pH 7.4, 150 mM NaCl, 0.5% (w/v) Sodium deoxycholate, 0.1% SDS, 1% (v/v) IGEPAL) supplemented with 1 mM PMSF protease inhibitor (Sigma) and 1x Protease Inhibitor Complete (Roche) prior to use and incubated for 30 min on ice followed by centrifugation at 13,000 rpm for 10 min at 4°C. Samples were separated on a 26-well 4-12% Bis-Tris SDS-PAGE gel (Life Technologies) and transferred onto a Hybond ECL (GE Healthcare) nitrocellulose membrane. The membrane was stained with Ponceau-S solution (Sigma, cat. #7170) for 5. Blocking with 5% (w/v) milk or Bovine Serum Albumine (BSA) in TBS-T (Tris Buffered Saline + 0.05% Tween-20) was followed by incubation with the primary antibody in blocking solution overnight at 4°C (see **Supp. Table 5** for antibodies). After washing 4× 5 min in TBS-T, the membrane was incubated with fluorescently-labelled secondary antibodies (IRDye; Licor) for 1 h at room temperature, washed 4x 5 min with TBS-T and scanned using the Licor Odyssey scanner.

### Annexin-V assay

HeLa cells were plated on 8-well LabTek II chambered coverglass (Nunc, cat.# 155409) and stained in Leibovitz (L-15) medium supplemented with 10% FCS and Pen/Strep, 100 μM Cisplatin (or 0.9% NaCl), FITC-labelled Annexin-V (Biolegend; 1:200), 7-AAD (7-amino-actinomycin D; Biolegend; 1:200), CaCl_2_ (2.5 mM; required for Annexin-V binding to phosphatidylserine) and TMRM (Tetramethylrhodamine, methyl ester; 20 nM). Imaging was performed using a Nikon Ti wide field microscope equipped with a sCMOS camera (Andor Zyla). Images of four channels (bright field; FITC, TRITC for TMRM; Cy5 for 7-AAD) were acquired every 15 min for 16 hours. Image analysis was performed manually using Fiji (ImageJ, NIH).

### Characterization of FRET pairs in fixed cells by imaging

The characterization of FRET pairs and sensors was done in fixed cells after transient transfection using Jetprime. 25 μM of the caspase inhibitors or DMSO (Dimethylsulfoxide) was added 2 hours before addition of 4 μM Staurosporine (Santa Cruz) or Ethyl Acetate (Sigma) as solvent control. Cells were fixed with 4% formaldehyde (Agar Scientific) for 10 min at room temperature. Imaging was performed with a multi-photon confocal laser scanning microscope (SP5, Leica Microsystems) tuned at 840 nm. Emission spectra were acquired with 10 nm detection bandwidth and step size. Fluorescence lifetime imaging was achieved with hybrid photo-multipliers tubes at the non-descanned port of the microscope. Time-correlated single-photon-counting was performed utilizing SPC-152 electronics by Becker&Hickl or a custom-build system utilizing time-to-digital concerted by Surface Concept GmbH.

### Live-cell time-lapse FLIM imaging

Two 8-well LabTek II chambered coverglass slides (Nunc) were prepared. On the control slide, parental HeLa cells were transiently transfected using JetPRIME with donors, the uncleavable controls and two acceptors (sREACh, tdNirFP), then imged by fast multi-colour FLIM (see **Supp. Notes 2** and **3**, and **Supp. Methods**) in imaging medium (Leibovitz (L-15) with 10% FCS and Pen/Strep). Time-lapse imaging was performed in the same conditions with HeLa-C239. Inhibitors and solvent controls were added 6 hours after transfection (25 μM for caspase inhibitors and LCL161 at 1 μM). Cisplatin at 100 μM, 0.9% NaCl and inhibitors at the same concentration used for pre-treatments when needed, was added in imaging medium after washing samples twice in PBS.

For double-thymidine block, cells were treated with 2 mM Thymidine (Sigma) at the time of plating and released in fresh medium 16 hour later. The same treatments were repeated after 8 hours. The second release was either synchronized with the time of imaging to image G1 cells or 7 hours before starting imaging to use G2 synchronized cells.

### Instrumentation and image-based biochemical analysis

Description and discussion of the instrumentation and image analysis developed for this work are described in **Supp. Notes 2** and **3**.

More detailed protocols, including a description of flow cytometry and ATP quantification are available in the **Supp. Methods**.

